# Pupil size reflects effort but not motor fatigability during a sustained motor task

**DOI:** 10.64898/2026.11.08.765154

**Authors:** Jenny Imhof, Caroline Heimhofer, Marc Bächinger, Sarah Nadine Meissner, Richard Ramsey, Nicole Wenderoth

## Abstract

Pupil size has been shown to reflect physical effort during brief, discrete motor tasks. It remains unknown whether pupil size similarly scales with effort during sustained repetitive movement, or whether it also reflects the accumulation of motor fatigability, a reduction in motor performance during prolonged movements. In our previous work, sustained maximal tapping induced motor slowing, a subtype of motor fatigability, characterized by decreasing movement speed and associated with increased antagonistic muscle coactivation. Next to these behavioural effects, pupil initially dilated and then gradually decreased over time. Whether this temporal change in pupil size is shaped by physical effort alone or also by motor fatigability remains unclear. Here, we combined a repetitive wrist tapping paradigm with force measurement, electromyography, and pupillometry, and paced participants at three different tapping speeds to dissociate physical effort from motor slowing, which has been shown to be speed-dependent. Fatigability-related motor slowing emerged selectively at the fastest speed. Moreover, we showed that antagonistic muscle coactivation increased more steeply at faster speeds, even in the absence of motor slowing, suggesting it may reflect an early change in motor control. While we found clear evidence that pupil size scaled with overall tapping speed, we did not find evidence that the temporal change of the pupil was pacing-dependent. These findings extend the pupil–effort relationship to sustained, dynamic motor tasks and suggest that pupil size is a marker of physical effort but not motor fatigability during sustained repetitive movement.

**Significance Statement:** Evidence linking pupil size to physical effort comes mainly from brief, discrete movements, yet many everyday activities require sustained movements. During such movements, performance can decline over time, a phenomenon called motor fatigability. Whether pupil size reflects only physical effort during sustained movements, or also the accumulation of motor fatigability, remains unknown. By pacing participants at different speeds, we manipulated both physical effort and motor fatigability, which emerges only at fast speeds. Pupil size scaled with physical effort, with little indication that it is additionally influenced by the accumulation of motor fatigability. We speculate that the fatigue-induced performance decline does not primarily arise because participants invest less effort over time, but rather because the motor system becomes progressively less efficient.

## Introduction

Pupil dilations that are not driven by changes in ambient light have been shown to reflect a variety of bodily states and processes such as arousal (Nieuwenhuis et al., 2011) and effort (Richer and Beatty, 1985; Hayashi et al., 2010; Zénon et al., 2014; Varazzani et al., 2015; van der Wel and van Steenbergen, 2018; Voudouris et al., 2023). The relationship between pupil and effort has been most extensively studied in the context of cognitive effort, with larger pupil dilations observed during more demanding cognitive control tasks and associated with better task performance (for a review: van der Wel & van Steenbergen, 2018). In contrast, relatively few publications have focused on the pupil’s link with physical effort (Zénon et al., 2014). Evidence linking pupil size to physical effort has mainly been derived from brief, discrete motor tasks, in which effort was operationalized as force magnitude, number of movements or movement amplitude (Richer and Beatty, 1985; Zénon et al., 2014; Varazzani et al., 2015; Voudouris et al., 2023). These studies reported strong phasic pupil dilations whose amplitude scaled with physical effort level. Whether this relationship extends to sustained movement is far less clear. To the best of our knowledge, the only study to address this question manipulated effort through cycling intensity and reported that pupil dilation increased with intensity (Hayashi et al., 2010). While these results suggest that pupil dilation is generally sensitive to physical demand, less is known about whether pupil size tracks effort during sustained movements over time.

Sustained fast, repetitive movements such as finger or wrist tapping are a reliable paradigm to induce motor fatigability in healthy individuals (Rodrigues et al., 2009; Bächinger et al., 2019; Heimhofer et al., 2024a, 2024b; Carro-Domínguez et al., 2025; Imhof et al., 2026a). Motor fatigability is defined as an objectively measurable decline in motor performance over time and should be distinguished from fatigue, the subjective feeling of tiredness or exertion (Kluger et al., 2013). While motor fatigability is a phenomenon we experience in everyday life, increased motor fatigability is a prominent symptom in many neurological disorders, such as Parkinson’s disease (Kluger et al., 2013) or multiple sclerosis (Kluger et al., 2013; Manjaly et al., 2019; Betti et al., 2026). This increased fatigability can impair the ability to maintain movements required for daily activities, such as walking (Van Geel et al., 2020). When induced by fast repetitive tapping, motor fatigability is characterised by a decrease in movement speed over time, referred to as motor slowing (Bächinger et al., 2019). Motor slowing has been associated with increased muscular activation and antagonistic coactivation per tap, indicating that, although participants slow down, each tap appears to require higher energetic costs (Bächinger et al., 2019; Imhof et al., 2026a). Together, these findings indicate that although tapping speed decreases during motor slowing, movement becomes more effortful as the motor system progressively requires more neural and muscular resources per movement.

In our previous work, we combined a sustained, effortful, maximal speed tapping task that induces fatigability-related motor slowing with pupillometry. We observed that pupil dilation first increased following movement onset, then gradually decreased as the participants’ tapping speed declined, before reaching a plateau (Imhof et al., 2026a). Similar pupil time-series patterns have been reported across other domains, such as cognitive and affective processes (Korn and Bach, 2016; Korn et al., 2017; Abivardi et al., 2023). However, the interpretation of this pupil time-series pattern in the context of motor fatigability remains unclear. Specifically, it raised the question of whether the observed decline in pupil size merely reflects the typical time course of an effort-evoked pupil response, or whether it also changes depending on the accumulation of motor fatigability.

Here, we address this question directly by testing whether the temporal change in pupil size varies with the accumulation of motor fatigability. We measure pupil size while systematically varying tapping speed such that faster initial movement speeds lead to pronounced motor slowing, while slower speeds produce little to no slowing. We hypothesized that pupil size scales with speed requirements of the tapping task. Additionally, we investigated whether speed-dependent differences in pupil size change over time as a function of whether motor fatigability accumulates.

## Methods

### Transparency and openness: pre-registration and open science statement

The study was preregistered prior to data collection (https://osf.io/rkavw). The preregistration included the research question, hypotheses, planned primary analyses, sample size, and exclusion criteria of this study. The preregistered primary outcomes were tapping speed and pupil size; all other outcome measures (tapping force and muscle (co)activation) were exploratory. In this paper, we report the preregistered sample size calculation, all data exclusions, all manipulations and all measures in the study. Throughout the manuscript, we highlight which analyses were preregistered and which ones were not.

### Participants

We tested 35 healthy participants (17 females; mean age = 24 ± 2 y). Participants were recruited via student mailing lists or online advertisements. All participants were between 18 and 35 years old, right-handed (assessed via the Edinburgh Handedness Inventory (Oldfield, 1971)), free of medication acting on the central nervous system, with no neurological and psychiatric disorders, and with normal or corrected-to-normal vision by contact lenses. Participants were asked to abstain from caffeine for at least four hours before testing. Prior to study participation, all participants provided written informed consent. The study was conducted with the approval of the research ethics committee of the canton of Zürich (KEK-ZH 2018-01078) and in accordance with the Declaration of Helsinki. Participation was compensated with 20 CHF per hour.

### Sample size estimation

The target sample size was determined prior to the data collection of this study using the pupil size data of five pilot participants. The pilot participants performed the same experimental task as tested in the study reported here. The participants were paced at three different tapping speeds (Fast, Medium, Slow; See *Behavioural Paradigm* in the *Methods* section for an explanation of the speed calculation). During each trial, participants had to tap at the paced target speed for 30s. The power analysis was conducted using G*Power. The pupil data was binned into 10s bins (i.e., 0-10s, 10-20s, 20-30s) and was treated as a 3 (*Time*: 0-10s, 10-20s, 20-30s) x 3 (*Pacing*: Fast, Medium, Slow) factorial design. The effect sizes for the differences between conditions within each bin were determined and then averaged over time. The smallest observed standardized mean difference was Cohen’s dz = 0.512 for the difference between the fast and medium pacing conditions. This effect size was chosen for the power analysis, yielding a sample size of 35 to achieve 90% power to detect this effect at an α = 0.05 significance level.

### General setup

The experimental setup was almost identical to the setup we used in a previous publication (Imhof et al., 2026a), in which we investigated pupil size dynamics during a wrist tapping task using the same combination of pupillometry and electromyographic (EMG) recordings. The experimental setup is depicted in Figure 1a (reproduced from Imhof et al., 2026a). The experiment was conducted in a quiet room with dim and constant ambient light. Participants were seated on a stable chair in front of a computer, with their heads against a chin and forehead rest to ensure a stable head position. The participant’s right hand was placed between two force sensors, with their forearm and elbow supported by an armrest. The forearm was fixated with Velcro as well as by a custom 3D printed lower arm fixation device to reduce compensatory movements from the forearm.

**Figure 1.**
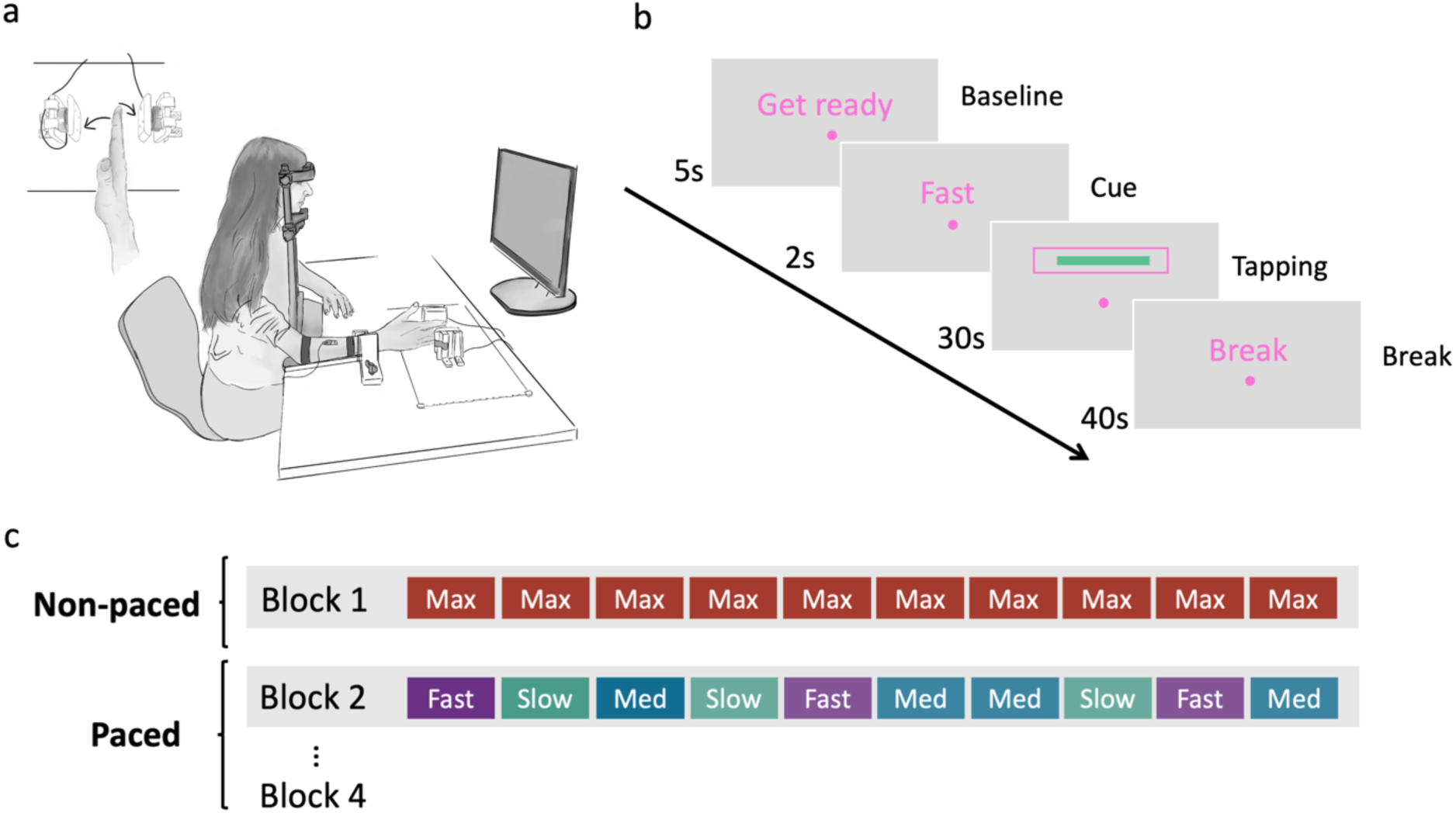
Experimental paradigm. (a) Experimental setup. Adapted from Imhof et al. (2026a). Participants were instructed to tap between two force sensors, so that they touched the sensors with their fingertips by alternating wrist flexion and extension. To reduce compensatory movements, the participants’ lower arm was fixated. (b) Visual display presented to participants during a paced trial. Each trial consisted of four phases: (1) a 5 s baseline period (“Get ready”), (2) a 2 s cue indicating the required tapping speed, (3) a 30 s tapping phase (pink target box), and (4) a 40 s rest period. During the tapping phase, participants received iso-luminant visual feedback on their tapping speed. A cursor moved up or down to reflect increasing or decreasing tapping speed relative to a target box representing the required target speed. The cursor turned from pink to green upon reaching the target. Trials of the non-pacing task followed the same structure besides that there was no visual feedback during the tapping phase. Instead of the visual feedback, “Go” appeared on the screen indicating that the participants should tap as quickly as possible during the tapping phase. (c) Overview of the experiment design. Each participant performed four blocks of wrist tapping. Block 1 consisted of 10 trials, during which participants were asked to tap at their maximal voluntary speed (non-paced task). Blocks 2 to 4 were paced. Each block of the pacing condition consisted of 10 trials. During each trial, participants were paced at a fast, medium or slow speed. Pacing conditions were presented in a pseudorandomized manner, in which each condition was presented at least once before a condition could be repeated.

### Behavioural paradigm

The behavioural paradigm consisted of two tasks, which were both performed by each participant: (1) a non-paced wrist tapping task at maximal voluntary speed, and (2) a paced wrist tapping task at three different target speeds relative to the participant’s individual speed in the non-paced task. To reduce potential biases in tapping behaviour due to strategic adjustments (e.g., tapping more slowly during the non-paced task to make the pacing task less demanding), participants were initially only informed about the non-pacing task. Only after completing the non-paced task did participants receive instructions about the paced wrist-tapping task. Data from the non-pacing task for one participant were lost due to a data-recording error. Consequently, all analyses of the non-pacing task were based on n = 34 participants, while analyses of the pacing task included all n = 35 participants.

#### (1) Non-paced wrist tapping task

Participants were asked to tap horizontally as quickly as possible for 30s, by alternately flexing and extending their wrist, so that their fingers touched two force sensors (Figure 1a). Each trial began with a baseline measurement, indicated by the message ‘GET READY’ (5s), during which participants mentally counted backwards from 100 in steps of four. This counting task was applied to minimize variability in pupil size due to fluctuations in attention or cognitive state (Ehlers et al., 2016). After the baseline, a cue (‘MAXI’) appeared (2s), signalling that participants could stop counting and that the following tapping phase required tapping at maximal voluntary speed. The subsequent ‘GO’ marked the start of the tapping phase (30s). Finally, ‘BREAK’ appeared, indicating a 40s rest period before the next trial began.

At the start of the experiment, participants completed a familiarization trial to ensure they understood the task correctly. During this familiarization period, the experimenter provided corrective and motivational feedback to promote accurate task performance (e.g., movement only from the wrist) and to encourage participants to perform the task at their maximal voluntary speed. After a short break, participants performed 10 experimental trials of the same task, during which the experimenter remained silent. Only participants who demonstrated more than 5% motor slowing during the non-paced wrist tapping task were included in the pacing task. Motor slowing was defined as the percentage decrease in tapping speed from the 0-10s bin to the 20-30s bin averaged across all non-paced trials:

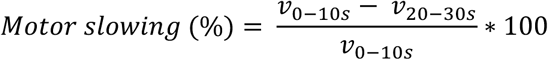

This check was implemented as participants were paced in the second task of the experiment based on their initial, medium, and final tapping speeds. Participants with less than 5% motor slowing would not have proceeded to the pacing task and would have been dismissed from the study. No participant had to be excluded based on this criterion.

#### (2) Paced wrist tapping task

In the second part of the experiment, participants performed a wrist tapping task while being paced at three different target speeds. The three target speeds were calculated using the data collected in the non-pacing task: The fast condition corresponded to the participants’ average tapping speed during the first 10 seconds of the non-pacing task, while the slow condition was based on the average tapping speed during the last 10 seconds. The medium condition represented the average speed of a gradual transition from fast to slow. Trials followed the same structure as the non-pacing task, with the following exceptions: instead of the ‘MAXI’ cue, a pacing cue (‘FAST’, ‘MEDI’, or ‘SLOW’) was presented between the baseline and tapping phase (Figure 1b). This cue informed participants of the target speed for the current trial. During the tapping phase, a cursor provided real-time feedback, moving up when participants were getting faster or down when participants were getting slower. A target box represented the desired target speed. When participants successfully tapped at the specified target speed, the cursor entered the target box and changed colour from pink to green. Participants were instructed to adjust their tapping speed based on the visual feedback so that the cursor moved to the target box and stayed inside the target box and green for as long as possible. The target box was always displayed at the same height for all three pacing conditions, to reduce potential gaze effects on pupil size. Throughout both parts of the task, participants were instructed to always fixate their gaze on a fixation dot to ensure that pupil size measurements were not confounded by eye movements.

In total, each condition was presented 10 times, resulting in 30 trials for the paced task. The 30 trials were split into 3 blocks with 10 trials each (Figure 1c). The conditions were presented in a pseudorandomized manner, so that each condition was presented once, before any condition could be repeated.

### Data acquisition

#### Tapping data acquisition

Taps and their force were recorded at 500 Hz using the two force sensors (KDEV-L-C1 10kg, Shenzhen Kaidechang Electronic Technology Co., Ltd., Shenzhen, China), whose signal was amplified (KDEV-B-0B, Shenzhen Kaidechang Electronic Technology Co., Ltd., Shenzhen, China), converted from an analogue signal to a digital signal using a Teensy (Teensy 3.2, PJRC.COM, LLC., Oregon, USA). The digitized force signal was transmitted via serial communication to the experimental computer, where it was read by the experiment script in MATLAB R2023a.

#### EMG data acquisition

We recorded the EMG signals of the musculus flexor carpi radialis (FCR) and the musculus extensor carpi radialis longus (ECR), the two main drivers of wrist flexion and extension, respectively. EMG signals were acquired using wireless surface electrodes (Tringo Wireless, Delsys, Natick, MA, USA), digitally sampled at 1000 Hz via a computer-controlled data acquisition device (CED Power 1401, Cambridge Electronic Design, Cambridge, UK), and band-pass filtered between 20 and 450 Hz. EMG activity and an analogue condition trigger from MATLAB were stored using Signal software (Cambridge Electronic Design, Cambridge, UK) for offline analysis.

#### Eye tracking and pupil data acquisition

Throughout the experiment, participants were instructed to focus on a fixation dot in the middle of the screen to ensure valid pupil size measurements not confounded by eye movements. Binocular pupil diameter and eye gaze were recorded at 60 Hz using the Tobii Pro Nano eye tracker (Tobii Technology) implemented in MATLAB R2023a using the Titta toolbox (Niehorster et al., 2020) in combination with the Tobii Pro SDK (version 1.10.1.2). At the beginning of each experimental block, the eye tracker was calibrated using a 9-point calibration procedure. To prevent luminance-driven changes in pupil size, all colours were isoluminant to the grey background (RGB 150 150 150), with relative luminance computed as a weighted linear combination of RGB components (Y = 0.2126R + 0.7152G + 0.0722B), reflecting the differential contribution of each colour channel to perceived brightness (https://www.w3.org/Graphics/Color/sRGB).

#### Stimulus presentation

The experiment was coded in MATLAB 2023a, and the stimuli presentation was controlled using the MATLAB-based software Psychtoolbox 3.0.19 (Brainard, 1997; Kleiner et al., 2007; Pelli, 1997).

### Data analysis

The pre-processing of the tapping, EMG and pupil data closely followed the pre-processing reported in (Imhof et al., 2026a). For completeness, we report the key steps below.

#### Behavioural data pre-processing

Before we could determine the tapping frequency (Hz) from the recorded force stream data, we first had to pre-process the raw data. Force signals from each sensor were offset corrected by subtracting the mean of the first two seconds (no tapping during this two seconds) and converted to Newton (N) using the following transfer function:

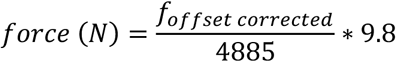

For each sensor, 2/3 of the root mean square (RMS) of the raw force signal during the tapping phase was used as a detection threshold, distinguishing genuine taps from noise and sensor crosstalk. Tap onset was identified when the signal exceeded this threshold with a minimum slope of 0.02 N/sample, subject to a minimum inter-tap interval of 100 ms and a maximum tap duration of 200 ms. Each detected onset was subsequently refined to the force peak within a 50 ms forward window. Automatically detected taps were then reviewed and manually corrected trial-by-trial using an interactive editor, allowing the experimenter to remove false positives and add missed taps by clicking directly on the force trace. Tap times and force peaks from both sensors were then merged and sorted chronologically. Tapping frequency was calculated as the inverse of the inter-tap interval (ITI), defined as the time elapsed between consecutive force events.

To account for missed taps (e.g., if participants did not properly hit the force sensor), identified when two consecutive events were registered by the same sensor, we applied a temporal interpolation procedure. The ITI of such events was compared to each participant’s mean movement cycle duration (defined as the time for one effector to complete a full movement cycle and return to its starting position). If the ITI fell within one standard deviation of the mean cycle duration, a single missed tap was inferred on the contralateral side and placed at the temporal midpoint between the two detected events via linear interpolation. ITIs outside this range (> mean ± 1 SD) were classified as artefacts; in these cases, the earlier tap was retained and the subsequent same-sensor tap was excluded. This procedure preserved the expected alternating tap structure while minimising distortions in cycle duration estimates. Interpolation rates were consistently low across all three pacing conditions (fast: 6.5% ± 0.7%; medium: 5.8% ± 0.8%; slow: 4.4% ± 0.7%).

#### EMG data pre-processing

The EMG data of the FCR and ECR muscles were pre-processed by first high-pass filtering the raw EMG signals at 20 Hz to remove low-frequency noise and motion artefacts (De Luca et al., 2010). The resulting signals were rectified and low-pass filtered at 20 Hz to extract the signal envelopes, which results in a smoothed representation of the muscle activity over time. The signal envelopes were then normalized to the global peak activity measured in each muscle. The area under the curve (AUC) of each normalized envelope was determined as a measure of overall muscle activation. To characterize the temporal evolution the antagonistic coactivation, we additionally computed a coactivation index for the FCR and ECR, defined as the overlapping area under the curve of the two normalized envelopes divided by the number of data points (Frost et al., 1997). Both the AUCs and the coactivation index were then normalized to tapping speed to account for differences in tapping speed across conditions and to isolate differences in muscle activation patterns not solely attributable to speed.

#### Pupil data pre-processing

Before pre-processing the pupil size data, we first visually inspected pupil size (in mm) and gaze to ensure that participants focused on the fixation dot throughout the trial. Trials with large eye movements or saccades (i.e., deviations >∼16° and ∼10° of visual angle on the x and y axes, respectively) affecting the pupil size were excluded. After visual inspection, pupil size data was pre-processed according to published guidelines using a standardized pipeline (Kret and Sjak-Shie, 2019). Invalid pupil diameter samples were identified and removed using a median absolute deviation (MAD) criterion, applied separately to dilation speed outliers (multiplier = 12) and large deviations from the trend line (multiplier = 30). Additionally, temporally isolated samples narrower than 50 ms and flanked by gaps exceeding 100 ms were excluded. Mean pupil size was then computed from the remaining valid samples across both eyes. At time points for which valid data was only available for one eye, the mean was estimated using the dynamic offset between the two eyes derived from time points at which both eyes provided valid measurements. The resulting time series was up-sampled to 1000 Hz and smoothed with a 4 Hz low-pass filter. Gaps exceeding 250 ms were left un-interpolated. After the pre-processing, we inspected all trials and removed those with more than 50% missing data during the baseline, cue presentation, or tapping phase. Overall, 44 of 1390 trials (3.2%) were excluded from the pupil analysis based on the visual inspection of raw and pre-processed data described above.

To increase statistical power by considering random fluctuations in pupil size over time (Mathôt et al., 2018), subtractive baseline correction was applied for each trial. Thus, the mean pupil size of the last second of the baseline phase before the start of the tapping phase was calculated and then subtracted from each data point of the tapping phase. To ensure that condition differences in the baseline-corrected pupil time series were not driven by differences in baseline pupil size, we averaged the uncorrected mean pupil size within the same baseline window for each participant and condition, and compared these values across conditions.

For an analysis requiring a continuous signal to identify time points at which the pupil time series differed between conditions, baseline-corrected time series were additionally subjected to cubic spline interpolation at the trial level prior to averaging across participants. Cubic spline interpolation was selected as it better captures the natural fluctuations in pupil size compared to linear interpolation (Fink et al., 2024).

Two summary measures were extracted from the baseline-corrected pupil time series: the initial peak pupil size, defined as the difference between baseline and the maximum value of the initial dilation (maximal pupil size in the first 5s), and the plateau, defined as the mean pupil size during the final time bin (20–30s). Both measures were derived from trial-averaged time series computed per participant and condition.

#### Statistical analysis

The primary analyses of this study, including analyses of the tapping speed and pupil size data, were preregistered. All deviations from the preregistration and exploratory analyses are clearly reported in the following subsections. Statistical analyses were conducted in R Studio (Version 4.5.2), unless otherwise specified, and the statistical significance level was set at α = 0.05.

To investigate how our outcome parameters, tapping frequency, tapping force, muscle activity and pupil size changed during the non-paced and the paced wrist tapping task, the data was binned into 10s intervals (0–10s, 10–20s, 20–30s) and averaged across trials. Linear mixed-effects models (LMEM) were fitted to the binned data using the lme4 package (Bates et al., 2015). The models of the non-pacing task included *Time* as a fixed effect, while the models for the pacing task included *Time, Pacing*, and their interaction. Adhering to the “keep it maximal” principles regarding random/varying effects (Barr et al., 2013), we started with models including random intercepts per participants and random slopes for all fixed effects and their interactions per participant.

When model convergence issues arose, we simplified the random effects structure of the models stepwise (For a full list with the final models applied see Supplementary Table 1). For each model, we assessed the distribution of residuals by comparing observed values to the expected uniform distribution using the DHARMa package (Hartig, 2024). Even though LMEM are quite robust, in cases where normality of residuals was violated (Kolmogorov-Smirnov test with p < 0.05), we obtained robust confidence intervals using bootstrapping with 1000 case resamples with the lmeresampler (Loy et al., 2025). This approach allowed us to validate statistical inference without relying on distributional assumptions. The resulting bootstrap confidence intervals were similar to model-based confidence intervals and supported the model findings, detailed comparisons are reported in the supplementary material (Supplementary Tables 2-5). We then generated Type III ANOVA tables using the lmerTest package (Kuznetsova et al., 2017). Partial eta squared (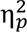) was calculated as an effect size for significant factors with the effectsize package (Ben-Shachar et al., 2020). Post-hoc analyses were conducted using the emmeans package (Lenth, 2024) and were adjusted for family-wise error rate (Bonferroni correction). The LMEM specifications for each outcome, together with analyses beyond LMEM, are described in the following subsections.

#### Data analysis of the non-pacing task

First, we analysed the data of the non-pacing task to determine whether we were able to replicate our previous findings regarding tapping speed, muscle (co)activation, and pupil size during a motor slowing task. Tapping force was additionally explored as a novel outcome measure. For each outcome parameter, LMEM with the fixed factor *Time* (i.e., 0-10s, 10-20s, 20-30s) and a random intercept per participant were used to test whether our outcome measures changed significantly over the course of the task. Muscle activity was modelled separately for the AUC of the FCR and ECR, and their coactivation index. Similarly, for the tapping force, separate models were fitted for the force recorded on the flexion and extension sides.

To test whether the expected decrease in pupil size correlated with motor slowing, a robust correlation was conducted using the MATLAB-based Robust Correlation Toolbox (Pernet et al., 2013). The decrease in pupil size was calculated as the difference between the maximum initial pupil dilation and the pupil size plateau, defined as the average pupil size of the last bin. Motor slowing was determined as the difference in tapping speed between the first and the last time bin. The Robust Correlation method was chosen as it protects against bivariate outliers (Pernet et al., 2013). No bivariate outliers were detected. Although multivariate normality was met as indicated by the Henze-Zirkler test, the data violated the assumption of homoscedasticity. We therefore applied Pearson correlation with significance assessed using bootstrapped 95%-confidence intervals, with effects considered significant if the interval excluded zero.

#### Data analysis of the pacing task

We analysed the data of the pacing task to investigate whether the change in tapping speed, tapping force, muscle (co)activation, and pupil size over time differed depending on the pacing condition. The same modelling approach as described above was applied to each outcome measure, with *Pacing* (fast, medium, slow), and *Time* (0–10s, 10–20s, 20–30s), and their interaction as fixed factors, and a random intercept per participant. For tapping speed and pupil size, the final model also included a random slope for the average effect of *Pacing* per participant. As before, separate models were fitted for the AUC of the FCR and ECR and their coactivation index, and for tapping force on the flexion and extension sides. A detailed list with the models fitted can be found in *Supplementary Table 1*.

The interaction between *Pacing* and *Time* did not reach significance in the LMEM on the binned pupil size data, but a non-significant result in this frequentist framework cannot be taken as evidence for the absence of an effect. We therefore performed the following additional, non-preregistered analyses to address whether pupil size indeed carries no information on the accumulation of fatigability over time. First, for each participant we computed the difference between the fast and slow baseline-corrected pupil time series and averaged it across participants to visually inspect how this difference evolved over time. We then binned this difference time series into 2.5s bins and fitted a LMEM with the factor *Time* and a random intercept per participant to the data, to test how the difference in pupil size changed over time. Second, we repeated the binned pupil size analysis at a higher temporal resolution of 2.5s as our 10s seemed too crude to capture the temporal pattern observed in the pupil time series. Finally, to formally quantify evidence for the null hypothesis, we subjected the binned pupil data to a Bayesian repeated measures ANOVA in JASP (Version 0.95.4) with the Pacing and Time as within-subject factors. We used JASP’s default Cauchy prior with scale r = 0.5. Inclusion Bayes factors were computed across matched models. Bayes factors were interpreted according to the classification scheme of Lee and Wagenmakers (2014), whereby BF values between 1–3 indicate anecdotal, 3–10 moderate, 10–30 strong, 30–100 very strong, and >100 extreme evidence.

In addition to the above mentioned LMEM with the binned pupil data, pupil size was subjected to three other preregistered analyses. First, a time series analysis was conducted subjecting the baseline-corrected pupil time series to the non-parametric alternative of a repeated measures ANOVA implemented in the MATLAB-based SPM1D toolbox for one-dimensional data (SPM1D M.0.4.8; https://spm1d.org/) (Pataky et al., 2013, 2015). While originally developed for the analysis of 1D kinematic, biomechanical, and force trajectories, SPM1D has also been applied to pupil size time-series (Meissner et al., 2024; Imhof et al., 2026a, 2026b). We conducted this analysis in addition to the binned LMEM to leverage their respective strengths: the LMEM provides a statistically powerful test of condition-related differences while accounting for inter-individual variability, whereas SPM1D preserves the full temporal structure of pupil dynamics, offering a detailed time-resolved characterisation of condition differences. Second, we analysed the initial increase in pupil size, defined as the difference between baseline and the maximum of the initial rise in pupil size, to test whether it scaled with the speed at which participants were paced. To test this, we used a LMEM with the fixed factor *Pacing* (fast, medium, slow) and a random intercept per participant. Third, we tested whether the plateau in pupil size towards the end of the tapping phase differed between conditions. Therefore, we used the average pupil size of the last time bin as the dependent variable and fitted a LMEM with the fixed factor *Pacing* (fast, medium, slow) and a random intercept per participant.

To ensure that condition differences in pupil size were not confounded by differences in absolute baseline pupil size, we compared the average pupil size during the final second of the baseline across pacing conditions as a control analysis using a LMEM with the factor *Pacing* and a random intercept per participant.

## Results

### Results of the non-pacing task

#### Wrist tapping at maximal speed led to progressively reduced motor output

We first assessed whether wrist tapping at maximal voluntary speed successfully induced motor slowing. Specifically, we were interested in whether tapping speed decreased during later time bins compared to the first bin. Indeed, tapping speed significantly decreased over time (LMEM: *F*(2,66) = 82.863, *p* < 0.0001, 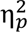 = 0.72, 95% CI for 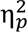: [0.62, 1.00]; Figure 2a), with significantly lower tapping frequencies in bin 10-20s and 20-30s compared to the first time bin (both *t* ≥ 8.302, *p* < 0.0001).

**Figure 2.**
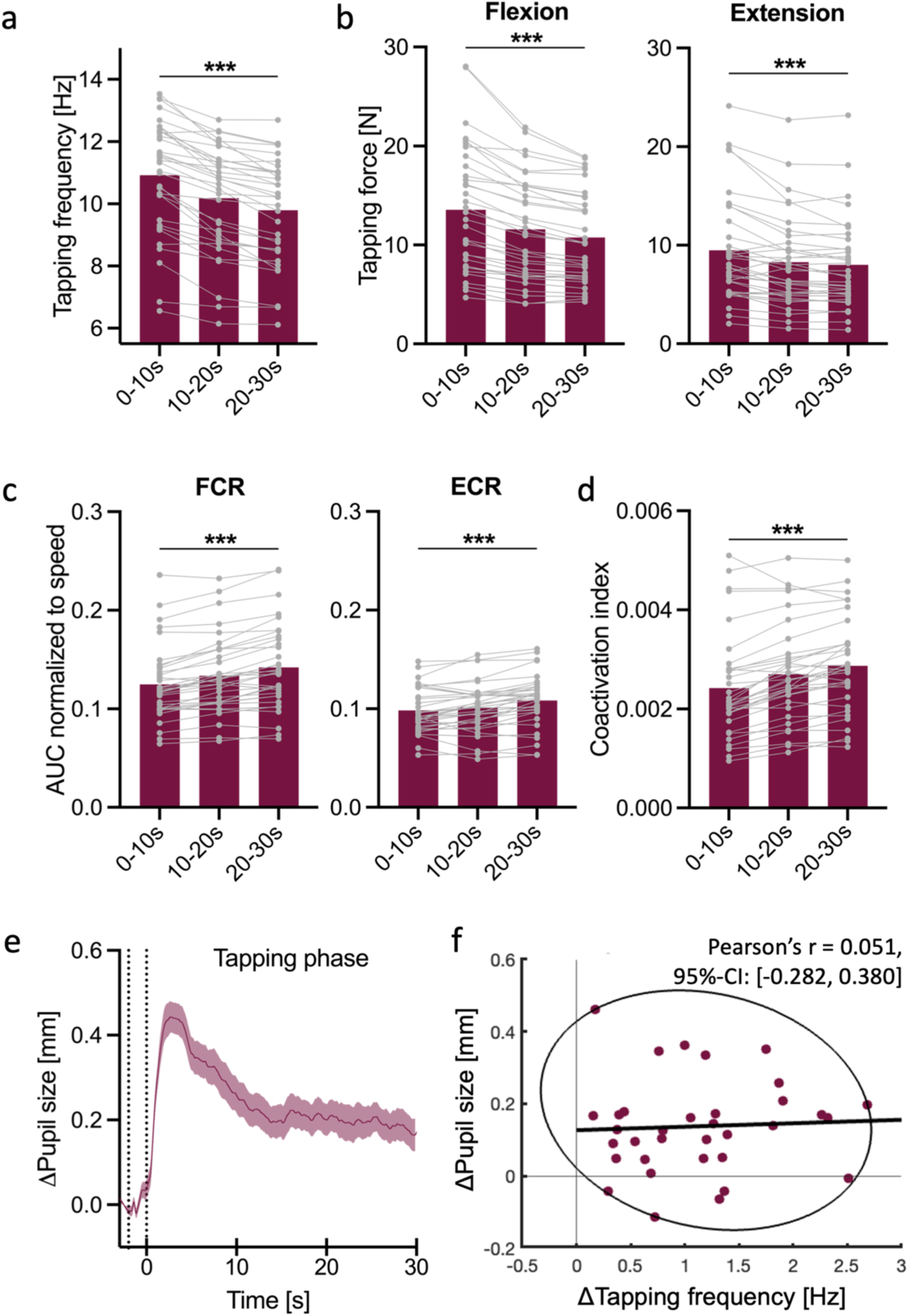
Non-paced wrist tapping at maximal voluntary speed. (a) Tapping frequency across time, (b) tapping force for flexion (left) and extension (right), (c) AUC of the flexor carpi radialis (FCR; left) and the extensor carpi radialis (ECR; right) electromyographic activity normalized to tapping speed, and (d) FCR–ECR coactivation index normalized to tapping speed; all binned, with group averages (red bars) and individual participant data shown in grey. (e) Baseline-corrected pupil diameter time series averaged across participants, showing the last second of the baseline phase, the two-second cue phase, and the 30s tapping phase. Vertical dotted lines mark the start of the cue phase and the end of the cue phase / start of the tapping phase. The shaded area represents the SEM. (f) Correlation between motor slowing and decrease in pupil size. Each dot represents one participant, and the ellipse denotes the data range included in the correlation. Asterisks denote significant pairwise differences between the first and last time bins: *p < 0.05, **p < 0.01, ***p < 0.001.

Having established that motor slowing occurred, we explored how the force per tap evolved over time. We found that both forces measured on the flexion and extension side significantly decreased over time (flexion side: *F*(2,66) = 57.800, *p* < 0.0001, 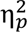 = 0.64, 95% CI for 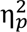: [0.52, 1.00]; extension side: *F*(2,66) = 22.019, *p* < 0.0001, 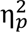 = 0.40, 95% CI for 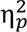: [0.24, 1.00]; Figure 2b), as tapping force was significantly reduced in the 10-20s and 20-30s time bins compared to the first time bin (flexion side: both *t* ≥ 7.433, *p* < 0.0001; extension side: both *t* ≥ 5.042, *p* < 0.0001).

These behavioural results indicate that tapping at maximal voluntary speed successfully induced motor slowing, with both tapping speed and force decreasing over time. Thus, motor output became progressively weaker during the trial.

#### Muscle activity and antagonistic coactivation significantly increased over time

We also investigated how muscle activation of the FCR and ECR, as well as their coactivation, changed over time during wrist tapping at maximal voluntary speed. All three outcome parameters significantly changed over *Time* (FCR: *F*(2,66) = 16.472, *p* < 0.0001, 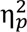 = 0.33, 95% CI for 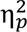: [0.18, 1.00]; ECR: *F*(2,66) = 50.079, *p* < 0.0001, 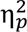 = 0.60, 95% CI for 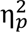: [0.48, 1.00]; coactivation index: *F*(2,66) = 36.404, *p* < 0.0001, 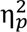 = 0.52, 95% CI for 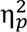: [0.38, 1.00]; Figure 2c and 2d). Muscle activity of the ECR and coactivation index per tap significantly increased across all time bins (ECR: all *t* ≤ -4.965, *p* < 0.0001; coactivation index: all *t* ≤ -3.197; *p* ≤ 0.006). FCR activation also increased overall (0-10s vs. 20-30s: t(66) = -5.601, p < .0001). However, this increase did not reach significance between the first two bins (0-10s vs. 10-20: t(66) = -1.716, p = 0.277).

Taken together, these findings show that during motor slowing, performing a wrist tap was associated with more muscle activation and antagonistic coactivation over time.

#### Pupil size initially increased and then decreased during motor slowing

We examined how pupil size changed over the course of the motor slowing task. Visual inspection of the baseline-corrected pupil time-series revealed that pupil first dilated and then gradually decreased over time, before it reached a plateau around 15s (Figure 2e). To investigate whether pupil size significantly decreased over time, we used the binned pupil size data for statistical analysis. A significant main effect of *Time* was revealed (*F*(2,66) = 32.295, *p* < 0.0001, 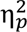 = 0.49, 95% CI for 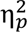: [0.35, 1.00]), with pupil size becoming significantly smaller during the 10-20s and the 20-30s bin compared to the 0-10s bin (both *t* ≥ 6.292, *p* < 0.0001).

#### Motor slowing and decrease in pupil size were not linked

Given that we observed a similar pattern in both tapping speed and pupil size, we wanted to test whether there is a correlation between the amount of motor slowing and the decrease in pupil size. The decrease in pupil size was defined as the difference between the maximum initial pupil dilation and the pupil size plateau, with the plateau calculated as the average pupil size of the last time bin. The conducted robust correlation revealed that there was no significant correlation between the two (Pearson’s r = 0.051, 95%-CI: [-0.282, 0.380]; Figure 2f). Therefore, the present data did not provide sufficient evidence for an association between these variables.

In summary, the results of the non-pacing task indicate that wrist tapping at maximal voluntary speed successfully induced motor fatigability, as reflected in a significant decrease in tapping speed and muscle force over time. Furthermore, we observed that tapping became more effortful, leading to higher muscle (co)activation per tap. Consistent with previous observations during motor slowing, pupil size initially increased before gradually decreasing over time. However, this decrease in pupil size did not significantly correlate with motor slowing.

### Results of the pacing task

#### Tapping speed indicates successful manipulation of physical effort and motor fatigability

We tested whether our pacing paradigm resulted in three distinct tapping speeds and whether motor slowing was most pronounced in the fast-pacing condition. Although the model’s residuals deviated from normality (Kolmogorov-Smirnov test, p < 0.05), bootstrap validation confirmed the robustness of our parameter estimates, producing confidence intervals that aligned with those derived from the model (see Methods, Statistical analysis). A detailed comparison of the model-based and bootstrap confidence intervals can be found in Supplementary Table 2. Tapping frequency differed significantly depending on pacing condition (LMEM main effect *Pacing*: *F*(2,34) = 97.535, *p* < 0.0001, 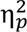 = 0.85, 95% CI for 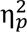: [0.77, 1.00]; Figure 3a), with the highest frequency in the fast condition across all time bins, followed by the medium and then the slow condition (all *t* ≥ 8.385, *p* ≤ 0.0001). The temporal change of the tapping frequency depended on the pacing condition (*Pacing x Time* interaction effect: *F*(4,204) = 22.651, *p* < 0.001, 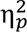 = 0.31, 95% CI for 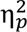: [0.22, 1.00]; Main effect of *Time* (*F*(2,204) = 5.734, *p* = 0.004, 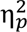 = 0.05, 95% CI for 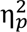: [0.01, 1.00]). As anticipated, we observed motor slowing as reflected by a significant decrease in movement speed from the initial time bin to the two consecutive ones only in the fast condition (both *t* = 6.414, *p* < 0.0001). In contrast, tapping speed remained stable in the medium condition (all *p* = 1.00) and even significantly increased in the slow condition from the first to the last time bin (*t* (204) = -4.197, *p* = 0.0001).

**Figure 3.**
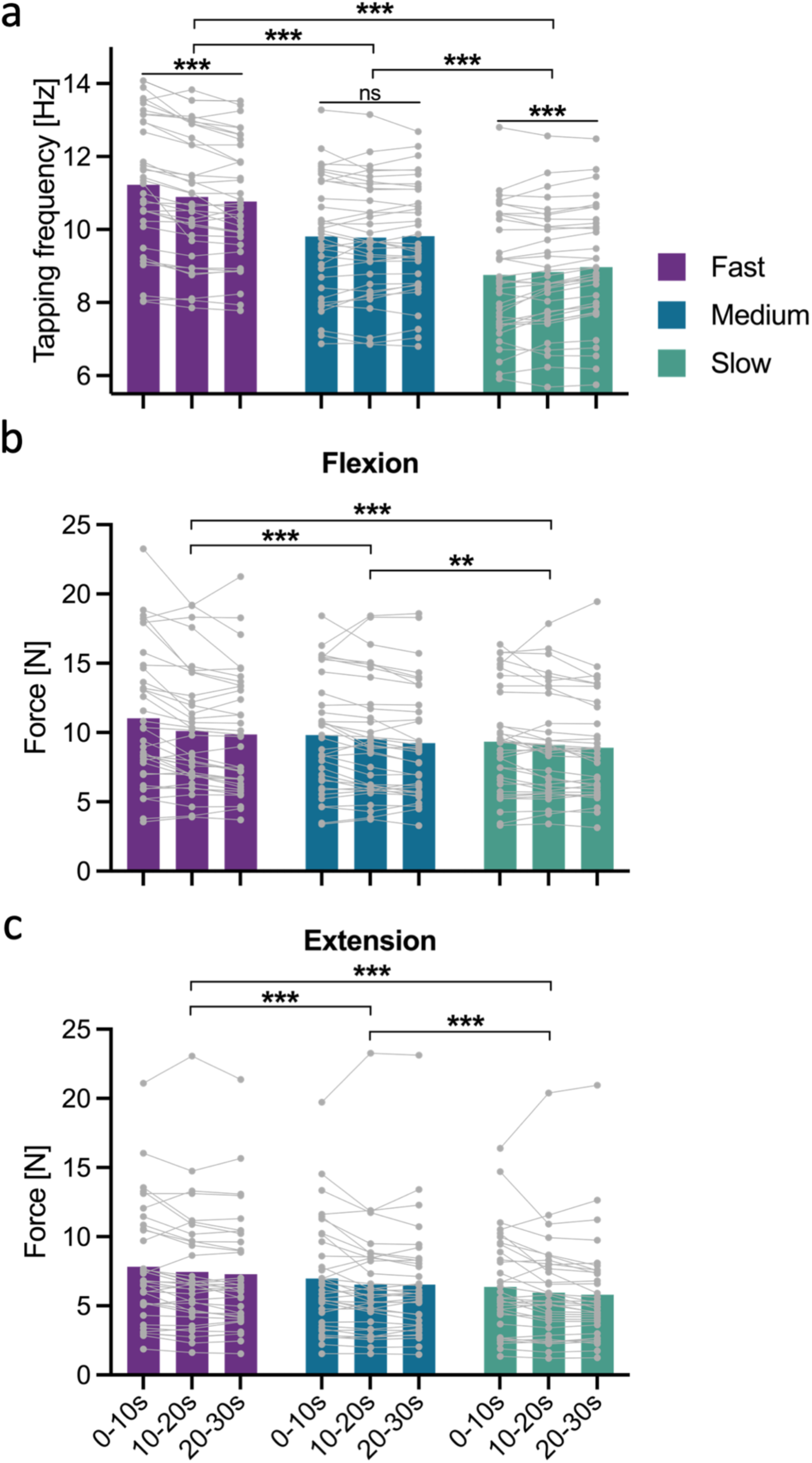
Behavioural data pacing task. (a) Tapping frequency (b) tapping force for flexion (left) and (c) extension (right) across pacing conditions. Binned data, with group averages as coloured bars (purple: fast; blue: medium; green: slow) and individual participant data shown in grey. Square brackets indicate comparisons between conditions, and horizontal lines indicate comparisons between time bins. Asterisks denote significant pairwise differences: *p < 0.05, **p < 0.01, ***p < 0.001.

These findings indicate that our pacing paradigm successfully established three distinct tapping speeds and selectively induced motor slowing only in the fast condition. This dissociation is critical, as it allowed us to disentangle whether subsequent outcome measures were driven solely by physical effort demands or additionally modulated by fatigability-related motor slowing.

#### Tapping force decreased over time

To characterize how tapping force evolved during our pacing task, we conducted an exploratory analysis for both the flexion and extension force. The residuals of both LMEM models violated normality (Kolmogorov-Smirnov test with *p* < 0.05). Thus, we applied bootstrap validation (see Methods Statistical analysis). The resulting confidence intervals broadly supported our model-based estimates, with the exception of the 10-20s : Medium and 20-30s : Medium interaction terms for the force on the flexion side, and for the force on the extension side, the 10-20s term. For these terms, the model-based CIs included zero while the bootstrap CIs did not (see Supplementary Table 3 and 4). Due to this discrepancy and the exploratory nature of the analysis, the findings reported below should be interpreted with caution.

The force per tap on the flexion side varied significantly across pacing condition (main effect of *Pacing*: *F*(2, 272) = 42.611, *p* < 0.0001, 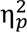 = 0.24, 95% CI for 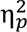: [0.17, 1.00]; Figure 3b) and declined over time (main effect of *Time*: *F*(2, 272) = 15.531, *p* < 0.0001, 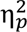 = 0.10, 95% CI for 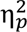: [0.05, 1.00]), with this decline not differing depending on conditions (interaction effect: *F*(4, 272) = 1.844, *p* = 0.121, 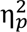 = 0.03, 95% CI for 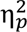: [0.00, 1.00]). Pooled post hoc tests indicated that flexion force per tap was the highest in the fast condition, followed by the medium and then the slow condition, with all pairwise differences reaching significance (*t* ≥ 3.027, *p* = 0.008). Across conditions, flexion force progressively decreased over time, with a significant reduction from the first to the second and first to the last time bin (both *t* ≥ 3.543, *p* ≤ 0.001).

The same pattern was observed for the force per tap on the extension side (Figure 3c). Significant main effects of *Time* (*F*(2, 272) = 12.259, *p* < 0.0001, 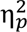 = 0.08, 95% CI for 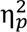: [0.03, 1.00]) and *Pacing*(*F*(2, 272) = 89.584, *p* < 0.0001, 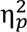 = 0. 40, 95% CI for 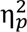: [0.32, 1.00]) were detected, while their interaction was not significant (*F*(4, 272) = 0.118, *p* = 0.976, 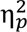 = 0.002, 95% CI for 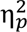: [0.00, 1.00]). The extension force applied differed significantly between conditions, with higher forces in faster conditions (all *t* ≥ 5.745, *p* < 0.0001). Tapping force declined significantly from the first to the second and from the first to the last bin (both *t* ≥ 3.716, *p* ≤ 0.0007).

Overall, these results suggest that participants were applying higher forces per tap when tapping at faster frequencies compared to slower frequencies. The force decreased over time with no significant differences between conditions.

#### Pacing modulates the time course of coactivation but not of overall muscle activity

We examined how muscle activity of the FCR and ECR, and their coactivation, changed over time during wrist tapping and whether this change depended on the pacing condition.

For the normalised AUC of the FCR, the muscle activity per tap differed between conditions (LMEM main effect of *Pacing*: *F*(2, 272) = 54.869, *p* < 0.0001, 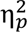 = 0.29, 95% CI for 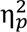: [0.21, 1.00]; Figure 4a) and changed over time (*Time* (*F*(2, 272) = 48.420, *p* < 0.0001, 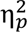 = 0.26, 95% CI for 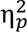: [0.19, 1.00]). The temporal change was not significantly influenced by pacing condition (*Pacing x Time: F*(4, 272) = 1.254, *p* = 0.288, 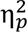 = 0.02, 95% CI for 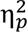: [0.00, 1.00]). Tapping at a faster pace was associated with higher overall muscle activity, and with higher muscle activity in the fast compared to the medium and slow pacing conditions (both *t* ≥ 7.703, *p* ≤ 0.0001). The muscle activity in the medium and the slow conditions did not differ significantly from each other (*t*(272) = 2.296, *p* = 0.067). FCR activity per tap increased progressively over time across all conditions (all *t* ≤ -3.559, *p* ≤ 0.001).

**Figure 4.**
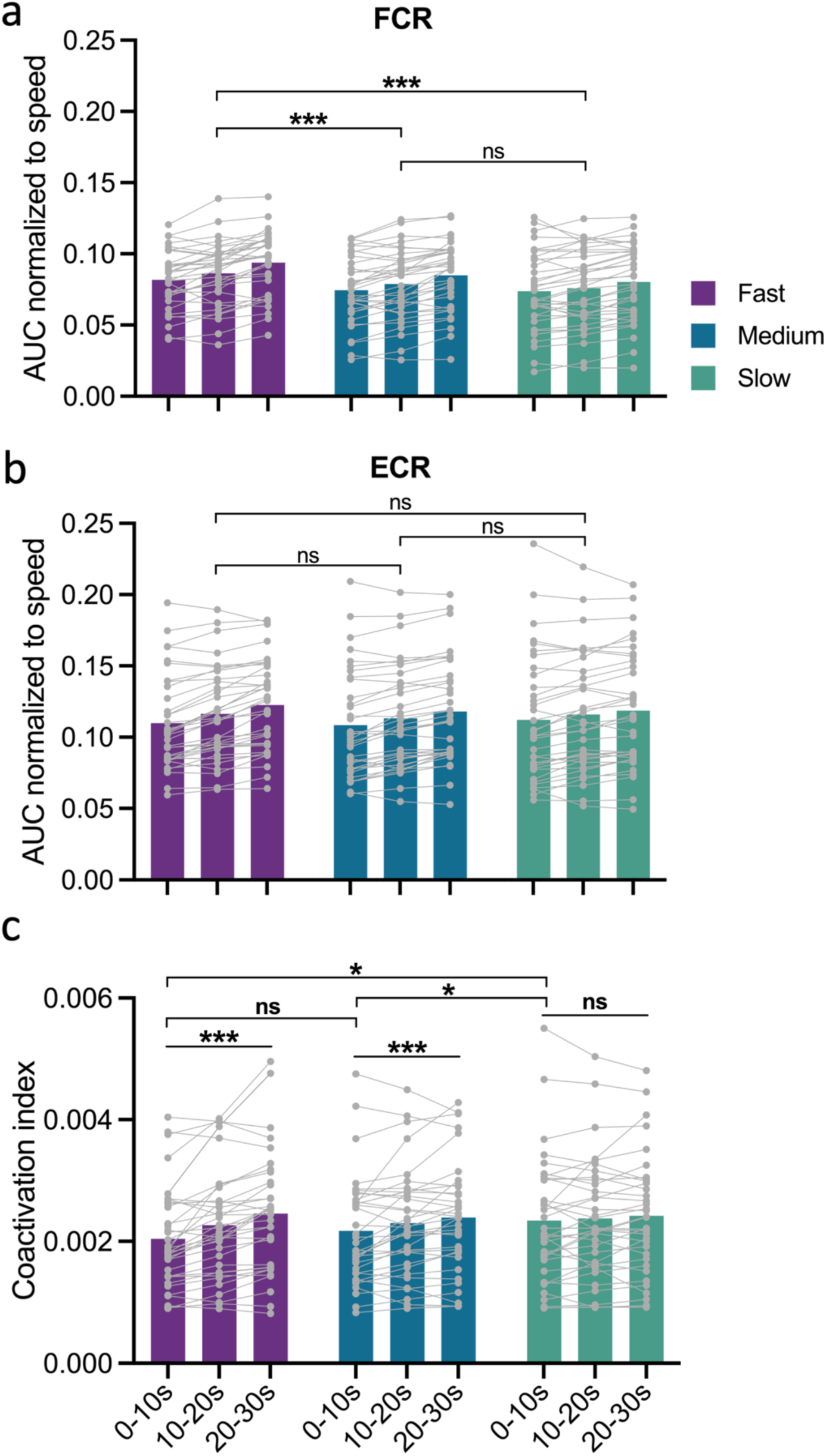
Muscle activity and coactivation during pacing task. AUC of (a) FCR (left) and (b) ECR (right) electromyographic activity normalized to tapping speed, and (c) FCR–ECR coactivation index normalized to tapping speed. All data binned, with group averages (purple, blue, and green bars) and individual participant data shown in grey. Square brackets indicate comparisons between conditions, and horizontal lines indicate comparisons between time bins. Asterisks denote significant pairwise differences: *p < 0.05, **p < 0.01, ***p < 0.001.

A similar pattern was observed for the normalised AUC of the ECR, with muscle activity increasing progressively over *Time* (*F*(2, 272) = 30.652, *p* < 0.0001, 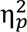 = 0.18, 95% CI for 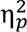: [0.12, 1.00]; Figure 4b). There was also a marginal main effect of *Pacing* (*F*(2, 272) = 3.102, *p* = 0.047, 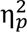 = 0.02, 95% CI for 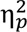: [0.00, 1.00]), whereas the interaction of *Pacing* x *Time* was not significant (*F*(4, 272) = 0.927, *p* = 0.448, 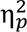 = 0.01, 95% CI for 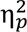: [0.00, 1.00]). Pooled post hoc tests revealed no significant differences between conditions (all *p* ≥ 0.095). ECR activity per tap increased significantly over time (all *t* ≤ -3.888, *p* ≤ 0.004). As the residuals of the model violated normality, we validated our estimates

using bootstrapped confidence intervals. The resulting bootstrapped confidence intervals were not fully consistent with our model-based results, as based on the bootstrapped intervals it seems that the interaction effect could be significant (see Supplementary Table 5). Given this discrepancy, these results should be interpreted with some caution.

We next examined how the coactivation index of the FCR and ECR changed during our pacing task. Coactivation was significantly modulated by pacing condition (*Pacing*: *F*(2, 272) = 4.146, *p* = 0.017, 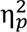 = 0.03, 95% CI for 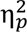: [0.00, 1.00]; Figure 4c), with significantly higher coactivation in the slow compared to the fast condition during the initial time bin (0-10s fast vs slow: *t*(272) = -3.884, *p* = 0.0004; all other are p ≥ 0.087). The coactivation index significantly changed over time depending on the pacing condition (*Time* x *Pacing*: *F*(4, 272) = 2.425, *p* = 0.048, 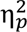 = 0.03, 95% CI for 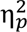: [0.00, 1.00]; Main effect of *Time* (*F*(2, 272) = 14.169, *p* < 0.0001, 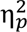 = 0.09, 95% CI for 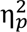: [0.04, 1.00]). Within the fast condition, coactivation significantly increased across all pairwise time comparisons (all *t* ≥ 2.466, *p* ≤ 0.043). Similarly, the coactivation index increased over time in the medium condition, with a significant increase from the first to the last time bin (0-10s vs. 20-30s: *t*(272) = −2.836, *p* = 0.015; other comparisons *p* ≥ 0.277). In contrast, no significant increase in coactivation was detected in the slow condition (all *p* ≥ 0.952).

Overall, our results indicate that muscle activity per tap increased over time for both the FCR and ECR, irrespective of pacing condition. Only the coactivation index of the two muscles showed an interaction effect between pacing and time. While at the beginning of the tapping the coactivation was highest in the slow condition, the coactivation significantly increased over time in the fast and medium condition but not in the slow one.

#### Pupil size scales with physical effort

To examine pupil size dynamics during our wrist tapping task at a high temporal resolution, we performed a time series analysis using the baseline-corrected pupil size time series. Visual inspection of the time series revealed a consistent pattern across conditions: pupil size first increased and then gradually decreased over time before reaching a plateau (Figure 5a). While the overall trajectory appeared qualitatively similar across conditions, pupil size was larger in the fast condition, followed by the medium condition, and then the slow condition. The statistical comparison supports the observation that pupil size differed depending on pacing condition (SPM{*z\**} = 6.726, 1 supra threshold cluster reaching from 2.2s to 33s with *p* = 0.001), with significantly larger pupil size in the fast compared to medium (SPM{t*} = 3.500, 3 supra threshold clusters with all cluster p = 0.001) and slow conditions (SPM{t*} = 3.318, 1 supra threshold cluster with p = 0.001). Pupil size during medium and slow conditions only differed during a few short temporal clusters (SPM{t*} = 3.645, 5 supra threshold clusters with the smallest cluster p = 0.006). Notably, condition differences were already present at the end of the cue presentation phase, with significantly larger pupil size in response to the fast cue compared to the slow cue, suggesting that anticipatory effort may have influenced pupil size.

**Figure 5.**
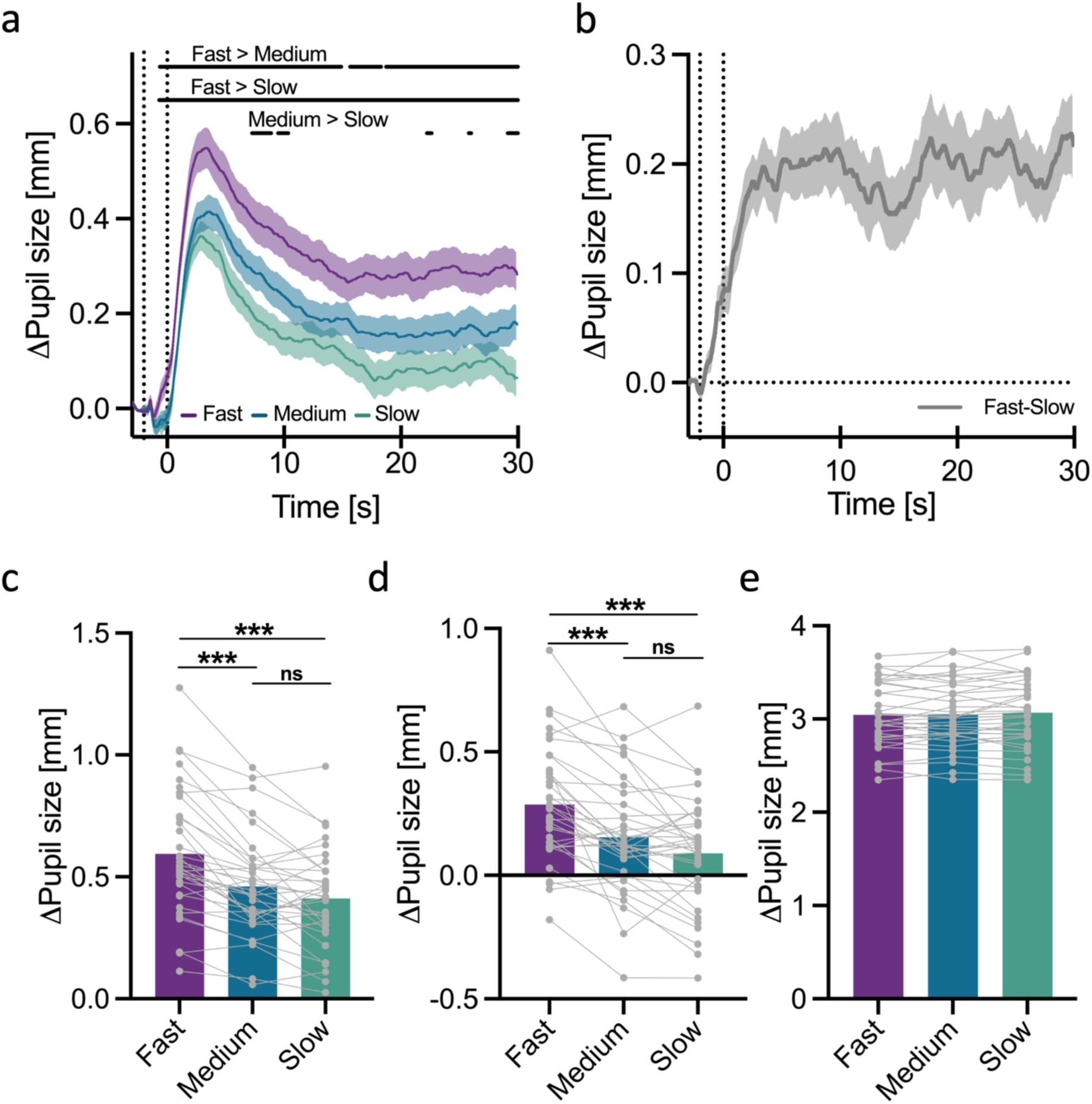
Pupil size during pacing task. (a) Baseline-corrected pupil diameter time series averaged across participants per condition, showing the last second of the baseline phase, the two-second cue phase, and the 30s tapping phase. (b) Average difference between the pupil size time series of the fast and slow condition. Vertical dotted lines in (a) and (b) mark the start of the cue phase and the end of the cue phase / start of the tapping phase. Shaded area represents the SEM. (c)The initial increase in pupil size per pacing condition. (d) The decrease in pupil size per pacing condition. (e) Absolute pupil diameter of the last second of the baseline phase per condition. In (c), (d) and (e) grey dots represent individual subject data, asterisks denote significant pairwise differences between conditions: *p < 0.05, **p < 0.01, ***p < 0.001.

While the time series analysis identified when pupil size differed between conditions, it does not permit the examination of how these differences evolved over time as a function of pacing condition. To address this, we conducted an analysis using binned pupil size data (10 s bins; Supplementary Figure 1a). This allowed us to directly test the interaction between pacing and time. This is informative with respect to fatigability, as only the fast condition was associated with motor slowing. Thus, if pupil size contains information on fatigability, we would expect a selective change over time in the fast condition but not in the other conditions, reflected in a significant *Pacing* x *Time* interaction. Our results showed that pupil size was modulated by both, *Pacing* (*F*(2, 34) = 23.884, *p* < 0.0001, 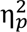 = 0.58, 95% CI for 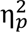: [0.39, 1.00]) and *Time* (*F*(2, 204) = 106.607, *p* < 0.0001, 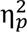 = 0.51, 95% CI for 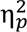: [0.43, 1.00]), but not by the interaction of these two factors (*F*(4, 204) = 0.168, *p* = 0.955, 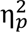 = 0.01, 95% CI for 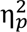: [0.00, 1.00]). As expected, pupil size was largest in the fast condition, intermediate in the medium condition, and smallest in the slow condition (all *t* ≥ 2.849, *p* ≤ 0.022). Collapsed across pacing conditions, pupil size significantly decreased over time, as pupil size was smaller in the middle and last time bin compared to the first bin (0-10s vs 10-20s and 0-10s vs 20-30s: *t* ≥ 11.982, *p* < 0.0001).

Since a non-significant effect of a frequentist analysis does not provide evidence on the absence of a true effect, we conducted the following additional analyses to further probe the non-significant *Pacing* x *Time* interaction. First, we visually inspected the average difference, and its variability, between the fast and slow baseline-corrected pupil time series (Figure 5b). This difference appeared to increase rapidly during cue presentation and the beginning of the tapping phase, after which it remained comparatively stable for the remainder of the tapping phase, suggesting that once the difference between conditions reached its peak, it stayed largely constant. We then tested whether this difference in pupil size changes over time, by fitting a LMEM to this difference data binned into 2.5s bins (See Supplementary Figure 1). The difference in pupil size changed significantly over time (*F*(11, 374) = 2.535, *p* = 0.004, 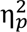 = 0.07, 95% CI for 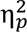: [0.01, 1.00]) with significant differences between the first time bin and bins 7.5-10s, 17.5-20s, 22.5-25s and 27.5-30s (all *t* ≤ -3.728, *p* ≤ 0.015), while all other comparisons did not differ significantly from each other. This pattern is consistent with the visual inspection, which showed a rapid increase in the difference at the very beginning of the tapping phase followed by comparatively stable values thereafter. We next asked whether the Pacing x Time interaction remained non-significant at a higher temporal resolution that better captures the temporal dynamics of our pupil size time series. Thus, we repeated the binned pupil-size analysis with 2.5 s bins. This higher-resolution analysis converged with our original result: pupil size was again modulated by both *Pacing* (*F*(2, 34) = 22.156, *p* < 0.0001, 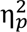 = 0.57, 95% CI for 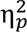: [0.36, 1.00]) and *Time* (*F*(11, 1122) = 78.434, *p* < 0.0001, 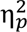 = 0.43, 95% CI for 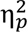: [0.40, 1.00]), but not by their interaction (*F*(22, 1122) = 0.722, *p* = 0.820, 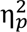 = 0.01, 95% CI for 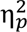: [0.00, 1.00]). While these results support the robustness of our finding, they do not directly address whether there is evidence in favour of the null hypothesis. To formally quantify evidence for the absence of a *Pacing* x *Time* interaction, we conducted a Bayesian repeated measures ANOVA on the 2.5 s binned data. This resulted in a Bayes Factor of BF_01_ = 2.280 in favour of the model excluding the interaction. Note that this is considered anecdotal evidence for the null hypothesis, i.e., that the temporal change of the pupil size does not depend on the pacing condition.

We preregistered two additional analyses focusing on two specific features of the pupil time series: the initial increase and the plateau at the end of the trial. As expected, both the initial increase in pupil size (Figure 5c) as well as the pupil size plateau (Figure 5d) differed significantly depending on pacing condition (initial increase: *F*(2, 68) = 34.642, *p* < 0.0001, 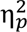 = 0.50, 95% CI for 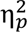: [0.36, 1.00]; plateau: *F*(2, 68) = 26.249, *p* < 0.0001, 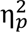 = 0.44, 95% CI for 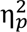: [0.28, 1.00]). During the initial rise, the fast condition differed significantly from both medium and slow (both *t* ≥ 5.899, *p* < 0.0001), while the medium and slow condition did not differ significantly from each other (*p* = 0.109). We detected the same pattern for the pupil plateau (20–30s), pupil was significantly larger in the fast condition compared to the medium and slow condition (both *t* ≥ 4.759, *p* < 0.0001), while the pupil size did not differ between the medium and the slow condition (*p* = 0.065).

To ensure that the condition differences observed in the baseline-corrected pupil size were not driven by differences in the absolute baseline pupil diameter, we tested whether the average pupil size during the last second of the baseline phase differed between conditions. The baseline pupil diameter did not differ significantly between conditions (*F*(2, 68) = 1.762, *p* = 0.315, 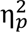 = 0.03, 95% CI for 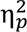: [0.00, 1.00]; Figure 5e), suggesting that changes in baseline corrected pupil size were not driven by large differences in baseline pupil diameter.

Together these results suggest that pupil size scaled with physical effort and that pupil size changed over time. However, we found no evidence to support the hypothesis that the gradual decrease in pupil size is affected by motor fatigability.

## Discussion

In this study, we used a pacing paradigm to manipulate physical effort and motor fatigability during repetitive wrist tapping while measuring tapping speed, tapping force, muscle activity, and pupil size. We showed that fatigability-related motor slowing emerged selectively at the fastest pacing speed, while tapping force declined across all conditions. Muscle activity and coactivation per tap scaled with pacing speed, but coactivation additionally showed a pacing-dependent temporal evolution, rising progressively in the fast and medium but not the slow condition. Pupil dilation scaled with tapping speed but showed no clear evidence for a pacing-dependent time course, suggesting that pupil size may primarily reflect the overall physical demand of the task rather than fatigability per se.

Our tapping speed results are in line with previous work, showing that tapping at fast speeds induced fatigability-related motor slowing (Rodrigues et al., 2009; Bächinger et al., 2019; Heimhofer et al., 2024a, 2024b; Carro-Domínguez et al., 2025; Imhof et al., 2026a). Moreover, we replicated the finding that motor slowing is speed-dependent and only appears if participants’ initial tapping speed is fast enough (Bächinger et al., 2019). Thus, pacing is a suitable tool to manipulate the fatigability state experimentally.

Having established that pacing successfully manipulated motor fatigability, we explored how tapping force changed over time in the context of motor slowing. Tapping force of both the flexion and extension decreased over time. To our knowledge, only one previous study has investigated tapping force during a fatigability-inducing repetitive tapping task. In this study, the opposite pattern was reported, namely a progressive increase in force over time, which was interpreted as a compensatory response to increasing perceived effort (Rönnefarth et al., 2019). These contrasting results may be explained by methodological differences: Rönnefarth et al. used a finger tapping task with the instruction to tap as fast and regularly as possible, which may have led to participants actively trying to maintain a consistent force output, whereas our wrist-tapping instructions emphasized speed alone. Furthermore, tapping with one’s fingers versus wrists involves smaller muscle groups and effectors, which could potentially lead to differences in force production. Rather than compensating by increasing force, our participants slowed down and produced less force while nonetheless recruiting more muscular (co)activation per tap, pointing toward rising movement cost per tap. Increased coactivation and muscle activity have been reported previously in the context of motor slowing (Bächinger et al., 2019; Imhof et al., 2026a). Even though the force trajectory differs between the two studies, both suggest that effort increases over the course of the task. Critically, this decline in force was not specific for the pacing condition showing motor slowing, suggesting that the gradual reduction in force over time is a general feature during sustained repetitive wrist tapping rather than a signature of fatigability-related motor slowing.

To better understand the muscular mechanisms underlying these behavioural changes, we next examined EMG activity. While the temporal increase in muscle activation per tap did not significantly change depending on pacing condition, antagonistic coactivation of the FCR and ECR muscle showed a pacing-condition-dependent time course. In the slow condition, coactivation was initially high and did not change significantly over time, whereas in the fast and medium conditions coactivation was initially low and then significantly increased over time. This lower initial coactivation in the fast and medium conditions, compared to the slow condition, suggests that reduced antagonistic muscle activity may be a prerequisite for achieving faster tapping speeds. The temporal increase in coactivation in the fast condition is consistent with previous work showing a significant rise in antagonistic coactivation during motor slowing (Bächinger et al., 2019; Imhof et al., 2026a). Furthermore, antagonistic coactivation has been linked to the release of cortical surround inhibition observed immediately after motor slowing, suggesting that such a loss in cortical inhibitory control manifests peripherally as changes in coactivation (Bächinger et al., 2019). Thus, the time-dependent increase in coactivation observed in our fast and medium conditions could be due to a decrease in cortical inhibitory control. Notably, coactivation increased even in the absence of measurable motor slowing in the medium pacing condition. This suggests that an increase in coactivation does not necessarily entail a decrease in motor performance, rather only once coactivation reaches a certain level does it lead to a significant decrease in tapping speed. Together, these results suggest that in order to tap at high speeds, a lower coactivation index is beneficial for efficient movement execution.

Finally, we investigated whether pupil size during sustained tapping tasks reflected physical effort or whether the gradual decrease in pupil size also contained information on the accumulation of motor fatigability. As expected, pupil size was the largest in the fast condition, followed by medium and then the slow condition. This finding aligns with previous research, in which pupil size has been shown to scale with the physical effort level generated (Richer and Beatty, 1985; Hayashi et al., 2010; Zénon et al., 2014; Varazzani et al., 2015; Voudouris et al., 2023) and extends it to a sustained tapping task. Moreover, we already see an increase in pupil size for the fast condition compared to the medium and slow conditions during the cue presentation phase, where participants were not yet tapping. This suggests that pupil size also reflects the anticipated effort, which has also been shown in effort-based decision-making tasks, especially when participants accept the physical effort rather than avoid it (Kurniawan et al., 2021). Together, these results indicate a general sensitivity of the pupil to physical demand. While in our study, pupil size scaled with overall tapping speed, its temporal change was not significantly modulated by pacing condition. Given that motor slowing appeared only in the fast condition, and that we did not find clear evidence for a significant interaction effect between time and pacing condition on pupil size, it is rather unlikely that average pupil size contains information about the accumulation of motor fatigability. Rather than reflecting a reduction in effort investment as performance declines, the temporal pattern of pupil size appears more consistent with the typical impulse-response dynamics of effort-evoked pupil responses, scaled with overall physical demand. This interpretation suggests that motor slowing does not arise because participants invest less effort over time, but rather because the motor system becomes progressively less efficient, resulting in decreasing motor output for the same invested resources.

While pupil dilations have been shown to reflect effort (Richer and Beatty, 1985; Hayashi et al., 2010; Zénon et al., 2014; Varazzani et al., 2015; van der Wel and van Steenbergen, 2018; Voudouris et al., 2023), it is important to acknowledge that pupil size is not an exclusive readout of effort. Grujic et al. (2024) describe pupil size as a one-dimensional readout combining multiple signals from multiple sources. Non-luminance related pupil size changes have been associated with different cognitive processes, such as decision-making (Urai et al., 2017) or reward processing (Schneider et al., 2018; Cole et al., 2022), ore more generally with different brain states that have been linked to arousal (Nieuwenhuis et al., 2011; Fink et al., 2024). Consistent with this, non-luminance-related pupil size changes have been linked to the activity of several neuromodulatory systems involved in regulating arousal, including the noradrenergic locus coeruleus (LC), the cholinergic basal forebrain, the serotonergic dorsal raphe nucleus, and orexin neurons of the lateral hypothalamus (Alnæs et al., 2014; Murphy et al., 2014; Joshi et al., 2016; Reimer et al., 2016; de Gee et al., 2017; Zerbi et al., 2019; Cazettes et al., 2021; Grujic et al., 2023). Importantly, arousal and effort are intertwined: the evaluation of task demands is thought to drive an increase in arousal, which in turn regulates the number of resources mobilized for the task at hand (Kahneman, 1973; Huang and Clewett, 2024). Evidence for the involvement of the LC-NA system in this process comes from research in monkeys: LC-NA activity during force production correlated with both the exerted force and the related pupil dilation (Varazzani et al., 2015), and pharmacological reduction of noradrenaline availability diminished force production (Jahn et al., 2018; Borderies et al., 2020). Thus, it is likely that pupil size as measured during our tapping task reflects physical effort and the associated arousal state.

Using a sustained wrist tapping paradigm combined with pupillometry, EMG, and force measurement, we show that tapping at fast speeds reliably induces motor slowing. By pacing participants at three different tapping speeds, we were able to dissociate physical effort from motor fatigability and demonstrate that pupil size scales with the overall physical effort of the task, whereas we did not find clear evidence that pupil size contains information about the accumulation of fatigability over time. This finding extends the pupil–effort relationship to sustained repetitive movement while suggesting that the temporal decline in pupil size reflects the typical dynamics of an effort-evoked response rather than a signature of motor fatigability. The pacing-dependent increase in antagonistic coactivation, which was more pronounced at faster speeds even in the absence of measurable motor slowing, further suggests that rising coactivation may reflect an early change in motor control that contributes to subsequent motor slowing. Together, these results advance our understanding of how the motor system becomes progressively inefficient during sustained movement and highlight pupil size as a marker of physical effort, with little indication that it also reflects motor fatigability, in sustained, dynamic motor tasks.

## Data availability statement

The data supporting the findings of this study will be openly available in the ETH Zurich Research Collection (link will be added upon acceptance of the manuscript).

## Author Contributions

J.I., C.H., M.B., S.N.M., R.R., and N.W. were involved in conceptualization and design of the study. J.I. programmed the task and analysis scripts and acquired the data. J.I., C.H., M.B., R.R., and N.W. planned the analysis. J.I. analysed the data. J.I., C.H., M.B., R.R., and N.W. interpreted the data. J.I. drafted the manuscript and all authors substantively revised it.

## Acknowledgments

The authors thank all participants for their study participation. Furthermore, we would like to thank Janis Senn for data collection, Jana Moos for assisting with the data analysis, Dr. Daniel Woolley for his technical support, and Dr. Snow (Xue) Zhang for her valuable contributions throughout the project. We would also like to thank Paige Howell for proof-reading, Eduardo Villar for the drawing of the experimental setup, and the Seminar for Statistics at ETH Zurich for statistical consulting. This work was supported by the Swiss National Science Foundation (SNSF) grant 32003B_207719. S.N.M is supported by the SNSF starting grant TMSGI1_226129. N.W. is supported by the National Research Foundation, Prime Minister’s Office, Singapore under its Campus for Research Excellence and Technological Enterprise (CREATE) programme.

