## Supplementary Material for "Pupil size reflects effort but not motor fatigability during a sustained motor task"

### Supplementary results

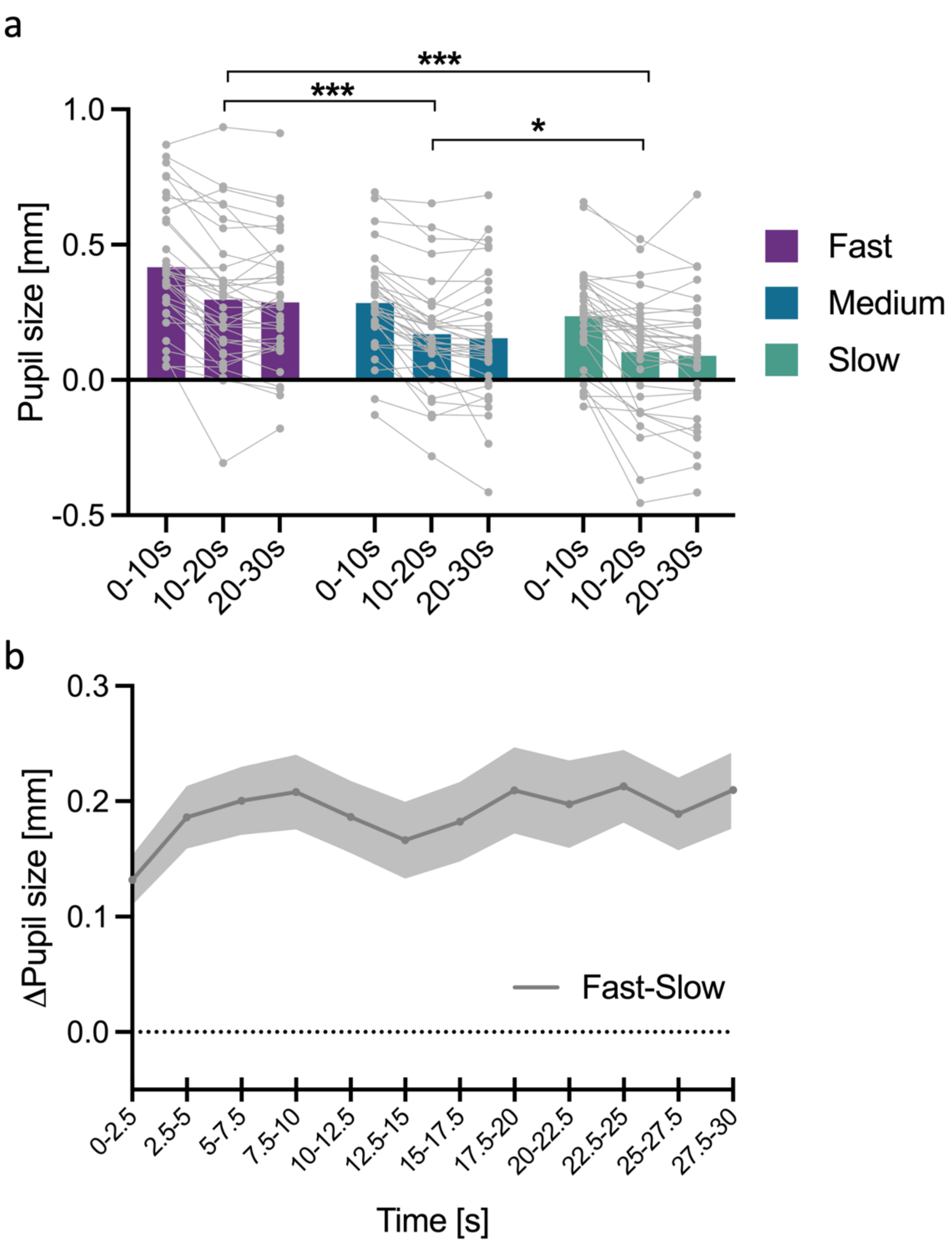


***Supplementary Figure 1. Pupil size during pacing task.*** (a) Baseline-corrected pupil diameter averaged across participants per time bin per condition (purple: fast; blue: medium, slow: green). These binned data was used for the statistical analysis reported in the main text (see Results: Pupil size scales with physical effort). Grey dots represent individual subject data, asterisks denote significant differences between conditions: *$p < 0.05$, **$p < 0.01$, ***$p < 0.001$. (b) Difference between the baseline-corrected pupil time series of the fast and slow pacing condition averaged across participants and binned into 2.5s time bins. Shaded area represents the standard error of the mean (SEM).

### Supplementary tables

**Supplementary Table 1. Overview of models fitted**. Always started with maximal model and reduced complexity stepwise. *Normality of residuals* shows whether the residuals met or violated normality (DHARMa, Kolmogorov–Smirnov test, $p < 0.05$ = violated). Bootstrap consistent shows, for models with violated normality, whether the bootstrap 95% CIs (1000 case resamples) agreed with the model-based 95% CIs (yes/no). A dash (–) means no bootstrapping was done because the residuals met normality. Detailed comparisons are given in Supplementary Tables 2–5

| Outcome measure | Model used | Normality of residuals | Bootstrap 95% CI |
|---|---|---|---|
| **Non-pacing part** | | | |
| Tapping speed | Speed ~ 1 + Time + (1 \| Participant) | Met | - |
| Tapping force flexion | Force_Flex ~ 1 + Time + (1 \| Participant) | Met | - |
| Tapping force extension | Force_Ext ~ 1 + Time + (1 \| Participant) | Met | - |
| AUC Flexor | AUC_Flex ~ 1 + Time + (1 \| Participant) | Met | - |
| AUC Extensor | AUC_Ext ~ 1 + Time + (1 \| Participant) | Met | - |
| Coactivation index | CI ~ 1 + Time + (1 \| Participant) | Met | - |
| Pupil size | Pupil ~ 1 + Time + (1 + Pacing \| Participant) | Met | - |
| **Pacing part** | | | |
| Tapping speed | Speed ~ 1 + Time * Pacing + (1 + Pacing \| Participant) | Violated | yes |
| Tapping force flexion | Force_Flex ~ 1 + Time * Pacing + (1 \| Participant) | Violated | no |
| Tapping force extension | Force_Ext ~ 1 + Time * Pacing + (1 \| Participant) | Violated | no |
| AUC Flexor | AUC_Flex ~ 1 + Time * Pacing + (1 \| Participant) | Met | - |
| AUC Extensor | AUC_Ext ~ 1 + Time * Pacing + (1 \| Participant) | Violated | no |
| Coactivation index | CI ~ 1 + Time * Pacing + (1 \| Participant) | Met | - |
| Pupil size | Pupil ~ 1 + Time * Pacing + (1 + Pacing \| Participant) | Met | - |
| Initial increase in pupil size | Pupil_Peak_mean ~ 1 + Pacing + (1 \| Participant) | Met | - |
| Pupil size plateau | Pupil_plateau_mean ~ 1 + Pacing + (1 \| Participant) | Met | - |

**Supplementary Table 2. Comparison of model-based and bootstrap confidence intervals for the tapping frequency**. Although the residuals of the LMEM violated normality, (Kolmogorov-Smirnov test with $p < 0.05$), bootstrap validation supported the robustness of our parameter estimates, yielding confidence intervals highly consistent with model-based estimates.

| Parameter | Estimate fixed effects | Model 95% CI | Bootstrap 95% CI |
|---|---|---|---|
| (Intercept) | 11.200 | [10.663, 11.788] | [10.8, 11.7] |
| 10-20s | -0.330 | [-0.431, -0.229] | [-0.437, -0.226] |
| 20-30s | -0.455 | [-0.556, -0.354] | [-0.611, -0.316] |
| Medium | -1.420 | [-1.602, -1.229] | [-1.57, -1.25] |
| Slow | -2.470 | [-2.786, -2.149] | [-2.73, -2.20] |
| 10-20s : Medium | 0.302 | [0.160, 0.445] | [0.191, 0.415] |
| 20-30s : Medium | 0.460 | [0.318, 0.603] | [0.312, 0.598] |
| 10-20s : Slow | 0.406 | [0.263, 0.548] | [0.298, 0.517] |
| 20-30s : Slow | 0.671 | [0.528, 0.814] | [0.508, 0.837] |

**Supplementary Table 3. Comparison of model-based and bootstrap confidence intervals for the tapping force on the flexion side**. Although the residuals of the LMEM violated normality, (Kolmogorov-Smirnov test with $p < 0.05$), bootstrap validation supported the robustness of our parameter estimates, yielding confidence intervals mostly consistent with model-based estimates.

| Parameter | Estimate fixed effects | Model 95% CI | Bootstrap 95% CI |
|---|---|---|---|
| (Intercept) | 11.033 | [9.667, 12.4] | [9.71, 12.2] |
| 10-20s | -0.916 | [-1.370, -0.462] | [-1.31, -0.509] |
| 20-30s | -1.173 | [-1.628, -0.719] | [-1.71, -0.597] |
| Medium | -1.219 | [-1.673, -0.765] | [-1.63, -0.85] |
| Slow | -1.682 | [-2.136, -1.228] | [-2.24, -1.16] |
| 10-20s : Medium | 0.639 | **[-0.003, 1.281]** | **[0.31, 0.965]** |
| 20-30s : Medium | 0.595 | **[-0.047, 1.237]** | **[0.256, 0.937]** |
| 10-20s : Slow | 0.689 | [0.047, 1.331] | [0.37, 0.996] |
| 20-30s : Slow | 0.720 | [0.078, 1.362] | [0.392, 1.08] |

**Supplementary Table 4. Comparison of model-based and bootstrap confidence intervals for the tapping force on the extension side**. Although the residuals of the LMEM violated normality, (Kolmogorov-Smirnov test with $p < 0.05$), bootstrap validation supported the robustness of our parameter estimates, yielding confidence intervals highly consistent with model-based estimates.

| Parameter | Estimate fixed effects | Model 95% CI | Bootstrap 95% CI |
|---|---|---|---|
| (Intercept) | 7.827 | [6.519, 9.135] | [6.77, 8.85] |

| | | | |
|---|---|---|---|
| 10-20s | -0.376 | **[-0.754, 0.002]** | **[-0.673, -0.052]** |
| 20-30s | -0.536 | [-0.914, -0.159] | [-0.85, -0.211] |
| Medium | -0.850 | [-1.227, -0.472] | [-1.10, -0.598] |
| Slow | -1.457 | [-1.834, -1.079] | [-1.86, -1.09] |
| 10-20s : Medium | -0.070 | [-0.604, 0.464] | [-0.305, 0.157] |
| 20-30s : Medium | 0.084 | [-0.450, 0.618] | [-0.244, 0.392] |
| 10-20s : Slow | -0.042 | [-0.576, 0.492] | [-3.44, 0.223] |
| 20-30s : Slow | -0.041 | [-0.575, 0.493] | [-0.442, 0.364] |

**Supplementary Table 5. Comparison of model-based and bootstrap confidence intervals for the AUC of the EMG envelope of the ECR**. Although the residuals of the LMEM violated normality, (Kolmogorov-Smirnov test with $p < 0.05$), bootstrap validation supported the robustness of our parameter estimates, yielding confidence intervals mostly consistent with model-based estimates.

| Parameter | Estimate fixed effects | Model 95% CI | Bootstrap 95% CI |
|---|---|---|---|
| (Intercept) | 0.11 | [0.098, 0.123] | [0.102, 0.119] |
| 10-20s | 0.007 | [0.002, 0.0113] | [0.005, 0.009] |
| 20-30s | 0.013 | [0.009, 0.018] | [0.01 0.017] |
| Medium | -0.001 | [-0.006, 0.003] | [-0.005, 0.002] |
| Slow | 0.003 | [-0.001, 0.008] | [-0.003, 0.009] |
| 10-20s : Medium | -0.002 | **[-0.008, 0.004]** | **[-0.004, -0.001]** |
| 20-30s : Medium | -0.003 | **[-0.010, 0.003]** | **[-0.006, -0.001]** |
| 10-20s : Slow | -0.003 | **[-0.010, 0.003]** | **[-0.006, -0.001]** |
| 20-30s : Slow | -0.006 | **[-0.0124, 0.001]** | **[-0.009, -0.004]** |
